# Behavioral heterogeneity in host-seeking and post-feeding suppression among disease vector mosquitoes

**DOI:** 10.1101/2025.06.18.660345

**Authors:** Takuya Uehara, Linhan Dong, Laura B. Duvall

## Abstract

Female mosquitoes seek out vertebrate hosts and consume their blood to obtain nutrients necessary for egg production, but host preference and host-seeking strategies differ markedly across species. These behaviors are also modulated by internal physiological states, such as the suppression of host-seeking after a full blood meal, a phenomenon that varies in timing and duration across mosquito species. We established a behavior monitoring and classification pipeline to systematically compare baseline host-seeking behavior and post-blood meal suppression in *Aedes, Anopheles*, and *Culex* mosquitoes. We found distinct behavioral signatures and notable interspecific differences in the onset and duration of host-seeking suppression. While *Aedes* and *Anopheles* host-seeking behaviors have been extensively studied in laboratory settings, comparable behavioral data for *Culex* have been limited, making direct comparisons across all three genera difficult. Our findings establish a unified behavioral framework for mosquito host-seeking across key vector species, providing insight into the ecological and physiological factors that shape host interaction and offering a foundation for improved modeling of disease transmission and vector control.

## Introduction

Female mosquitoes’ strong attraction to vertebrate hosts makes them the world’s deadliest animals. While thousands of mosquito species exist globally, only a subset pose significant public health threats due to their preference for human hosts and their ability to transmit pathogens responsible for diseases such as dengue, malaria, and West Nile fever.^1^ Females rely on a combination of sensory cues including carbon dioxide (CO_2_), host odor, and body heat to locate and bite hosts.^2^ However, mosquito species vary in host preference (human vs. non-human) and the timing of their activity (diurnal vs. nocturnal).^3^ *Aedes* species are typically daytime biters, while *Anopheles* and *Culex* primarily bite at night.^4–8^ In addition to external cues, host-seeking behavior is tightly regulated by internal physiological state. Across multiple mosquito genera, newly-eclosed females are not immediately receptive to host cues and must mature before they can blood feed.^9^ After consuming a full blood meal, females enter a transient period of host-seeking suppression, during which their responsiveness to host cues is reduced until eggs are laid.^2,10,11^ This phenomenon has been best characterized in *Aedes aegypti*, and while both *Aedes* and *Anopheles* species have been well-studied in laboratory settings, comparable behavioral data for *Culex* remain limited. Host-seeking suppression is thought to involve mechanosensory and reproductive signals, but the relative contributions of these factors remain poorly understood across species, in part due to a lack of comparative data.

Typically, a single blood meal is sufficient to initiate and complete a gonotrophic cycle (egg development and oviposition), but multiple feeding events have been reported, particularly in field settings.^12–15^ In *Anopheles gambiae*, some females undergo a “pre-gravid” phase, requiring two blood meals on different nights to complete the first gonotrophic cycle.^16^ Larval nutrition can also influence adult female host-seeking behavior; *Aedes aegypti* females with insufficient larval nutrients are more likely to require multiple blood meals to support egg production.^17^ Even among laboratory studies, host-seeking dynamics differ between species. A comparative study of several *Aedes* and *Anopheles* species found that *Anopheles gambiae* females resumed host-seeking within 24 hours after a blood meal, prior to oviposition, unlike *Aedes aegypti* and *Aedes albopictus* females, which typically show longer periods of host-seeking suppression.^13^ However, other research suggests that host-seeking behavior in blood-fed *Anopheles gambiae* is suppressed for approximately 40 hours post-feeding, until oocytes mature or eggs are laid.^18^ While most research has focused on *Aedes* and *Anopheles, Culex* mosquitoes also feed on humans and can transmit pathogens such as West Nile virus.^19,20^ Although seasonal modulation of host-seeking occurs in some *Culex* species, laboratory studies specifically addressing postprandial host-seeking suppression in *Culex* are lacking.^21^ Together, these findings underscore the need for a unified, comparative framework to understand how host-seeking behavior is regulated across mosquito genera.

Immediately following blood consumption, distention of the abdomen is believed to contribute to short-term suppression of host-seeking via mechanosensory signals. Female mosquitoes reliably ingest blood meals that double their body weight and classic studies have shown that the act of blood-feeding can be experimentally uncoupled from the nutritional content of the blood. These studies identified ATP as a key phagostimulant that triggers engorgement when presented with sodium chloride and sodium bicarbonate.^22^ In *Aedes aegypti*, abdominal distension alone, induced by ingestion of non-nutritive saline, is sufficient to inhibit host-seeking^23^ and the degree of distension, determined by meal volume, influences the duration of suppression.^24^ However, the role of distension signals appears to vary across species; in *Culex tarsalis*, a small proportion of females captured while seeking avian hosts had visible blood in their midguts, suggesting that abdominal distension may play a less prominent or differential role in suppressing host-seeking in this species.^15^ In *Aedes*, abdominal distension not only suppresses host-seeking but can also promote egg development, engaging downstream reproductive pathways that may further contribute to behavioral suppression.^25,26^ After distension wears off, neuropeptide signals regulate host-seeking in *Aedes aegypti* and *Anopheles stephensi*, although their anatomical localization and downstream targets of action are likely distinct.^27–30^ Reproductive signals, including ecdysone and vitellogenin, have also been shown to impact host-seeking behavior. Ecdysone release initiated after a blood meal suppresses host-seeking in *Anopheles freeborni*.^31^ In *Aedes albopictus*, sugar feeding alone is reported to be sufficient to drive expression of vitellogenin in the fat body and suppress host-seeking.^32^

Host-seeking is typically measured using endpoint assays that assess attraction to traps baited with host cues.^33–35^ However, females often show increased flight activity before host-seeking fully resumes, potentially influenced by egg-laying behaviors that also elevate activity levels in the absence of active host-seeking.^36,37^ Advances in ethology increasingly use machine learning pipelines to analyze animal behavior, and these tools can be applied to mosquitoes to characterize behavioral modalities.^38^ By generating detailed postural readouts of host-seeking and its suppression after blood-feeding, these approaches can provide deeper insight into behavioral states beyond general activity levels.^40,41^ Here, we established an automated recording platform and a behavioral classification pipeline to systematically compare the postural responses to human host cues across mosquito species and examine how this changes after blood feeding. While prior research has identified key signaling pathways that regulate host-seeking suppression, our findings reveal interspecific behavioral variation, which suggests species-specific expression or modulation of regulatory processes. These findings will inform future experiments intended to manipulate candidate signaling pathways to understand the mechanistic basis of these differences.

## Results

### Multipoint tracking identifies differential behavioral responses to human odor and CO_2_

To assess mosquito behavioral responses to host cues, we designed an assay to record high-resolution videos of individual females in controlled arenas following a pulse of CO_2_ and human odor (Figure 1A). Each 9-minute trial includes 1 minute of baseline activity recording, followed by a 3-minute exposure to 10% CO_2_ and human odor, followed by 5 minutes of post-stimulus recording (Figure 1B). To investigate host-seeking behavior across evolutionary lineages, we tested six mosquito species representing three major genera: *Aedes aegypti* and *Aedes albopictus* (*Aedes*), *Culex quinquefasciatus* and *Culex tarsalis* (*Culex*), and *Anopheles stephensi* and *Anopheles gambiae* (*Anopheles*). This selection allowed us to compare patterns of host-seeking regulation both within and across genera as *Aedes aegypti* and *Aedes albopictus* diverged ~30– 50 million years ago (MYA), *Culex quinquefasciatus* and *Culex tarsalis* diverged ~10–20 MYA, and *Anopheles gambiae* and *Anopheles stephensi* diverged ~35–55 MYA; the *Aedes, Anopheles*, and *Culex* genera themselves are estimated to have diverged from one another over 150 million years ago.^42–44^ We aimed to identify behavioral features that are differentially expressed and to evaluate whether these differences correspond to evolutionary relationships among the species.

**Figure 1:**
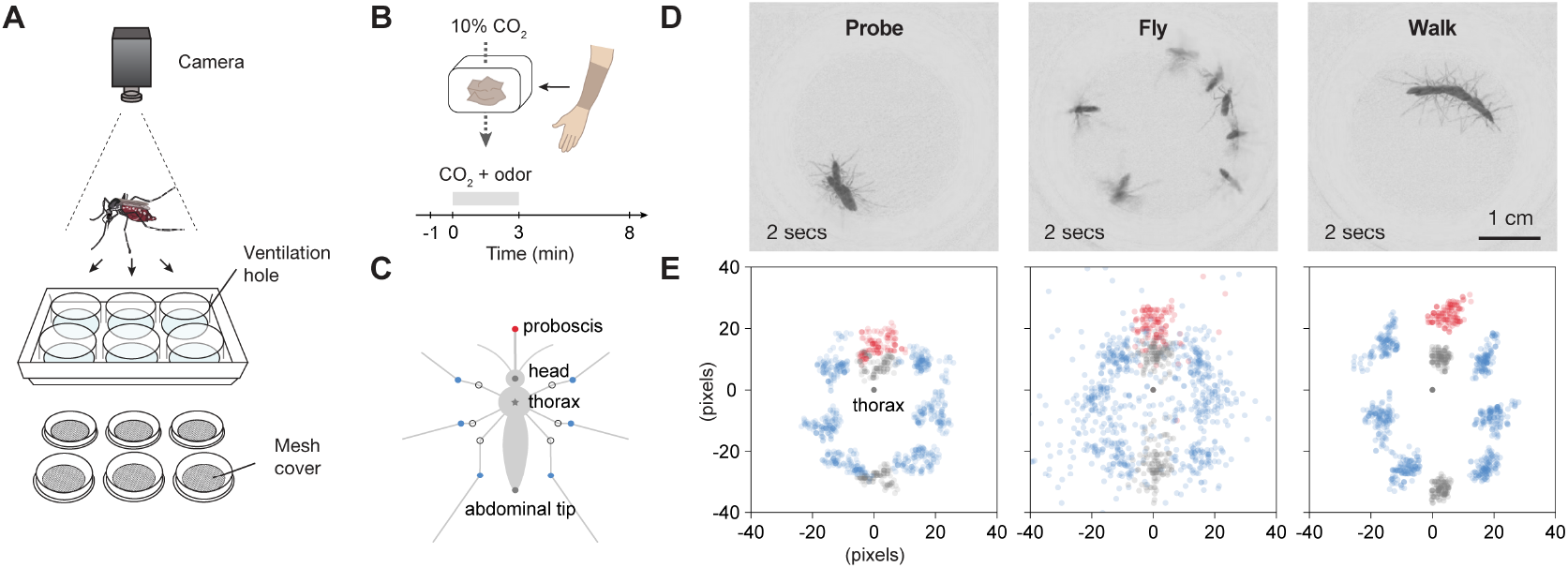
High resolution tracking of behavioral responses to human host cues. (A) Schematic of the behavioral arena. Six females were individually housed in the wells of a standard 6-well tissue culture plate with adaptations to allow for stimulus delivery and ventilation. (B) Time course of stimulus delivery and recording. (C) Schematic of track point placement across tested species. (D) Two-second time projection of example behaviors in *Aedes aegypti*. Every 20 frames (0.33 sec intervals) in the 2-second recording fragment were used to generate a minimum projection for visual clarity. (E) Track point distribution of respective two-second behavioral fragments. Track point coordinates were transformed using thorax as origin and head-abdominal tip as y-axis direction.

To generate a detailed representation of mosquito movement, we trained a neural network using DeepLabCut^45^ to label 16 postural key points on multiple *Aedes aegypti* and *Aedes albopictus* mosquitoes (Figure 1C). The resulting model demonstrated reliable tracking of animal key points in *Aedes* and *Culex* species tested (Figure S1A), but showed compromised performance when tracking *Anopheles gambiae*, likely due to differences in body shape and size. Therefore, we trained a separate model for *Anopheles* species that improved tracking performance (Figure S1B). This method captures fine-scale postural changes, enabling high precision analysis of movement dynamics. Using multipoint tracking data, we performed machine learning-based behavior clustering with Variational Animal Motion Encoding (VAME).^46^ We trained this unbiased behavioral clustering algorithm using data from the component species of each genus, instructing it to classify 30 distinct postures per genus. This approach reliably resulted in the identification of 6–10 key postures related to walking, flying, and probing. Since the key probing pose is often identified as a transient “bite”, we further processed the posture data to recognize signature “probe-walk-probe” postural combinations to accurately report a prolonged probing behavioral state, resulting in a pipeline that was effectively able to distinguish flying, walking, and probing behaviors (Figure 1D and E). The remaining postures included stationary states or subtle leg movements, such as vertical movements of the hindlegs and stroking the proboscis with the forelegs, likely reflecting rest and grooming behaviors. Manual behavioral classification on a subset of activity confirmed that the trained classifier showed high agreement with human-labeled behavior (96.0 ± 1.2% accuracy in *Aedes*, 92.8 ± 1.4% in *Anopheles*, 94.4 ± 1.3% in *Culex*) (Figure S1C).

**Figure S1.**
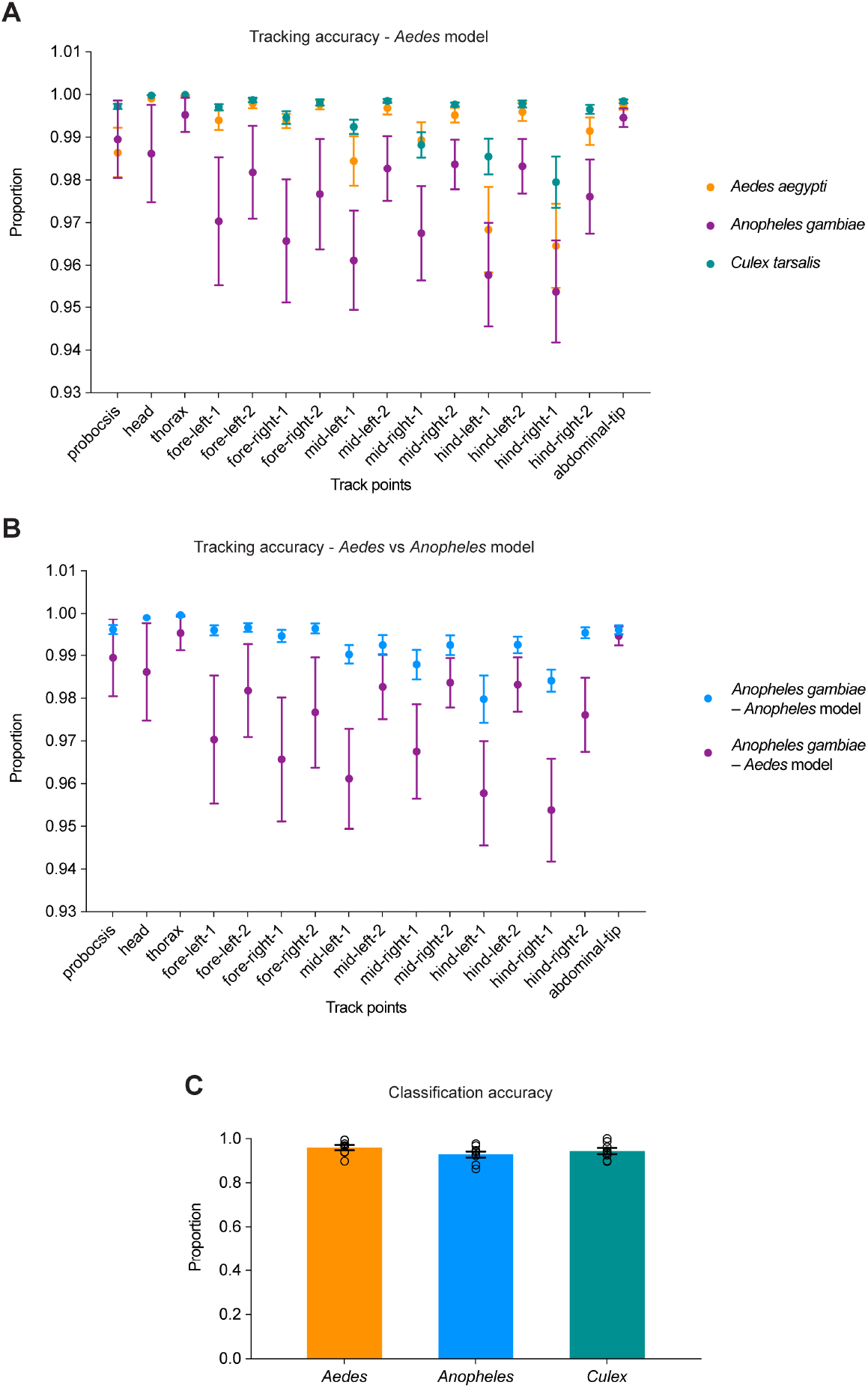
Performance metrics of pose estimation and classification. (A) Average confidence score of each key point using the *Aedes* model (values range from 0 to 1, indicating prediction reliability, N = 8 females). Mean and SEM. (B) Comparison of average confidence scores obtained from the original *Aedes* and the new *Anopheles* models in *Anopheles gambiae* (N = 8 females). Mean and SEM. (C) Example pose classification accuracy between manual annotation and the trained classifier (VAME) (N = 8 females). Mean and SEM.

### Host-seeking postural repertoire varies between mosquito species and genus

Using automated delivery of a combination of carbon dioxide and human odor (Figure 1B), we recorded behavioral responses of all six species tested at their preferred host-seeking time of day, daytime in *Aedes* and nighttime for *Anopheles* and *Culex* species (Figure 2A). All species tested exhibited a post-stimulus increase in activity (Figure 2B), characterized by enhanced walking, probing, and flight behaviors that began after stimulus onset and continued beyond its offset (Figures 2C and D and Figure S2). We present data from one species per genus in Figure 2, with additional species included in Figure S2. Notably, *Aedes albopictus* females displayed variable responsiveness (Figure S2), while *Culex tarsalis* and *Culex quinquefasciatus* showed a strong preference for walking in this assay (Figure 2E). Overall, *Culex* mosquitoes were more behaviorally active than *Aedes* or *Anopheles* species, spending a greater proportion of time walking and probing (Figure 2E). To globally compare behavioral differences, we performed a Principal Component Analysis (PCA), which showed that species within the same genus occupied similar PCA space, suggesting that host response behavioral repertoires are more conserved within genera (Figure 2F). These findings confirm that all species respond to host cues and also reveal distinct patterns of movement and posture that may reflect distinct host-seeking strategies.

**Figure 2:**
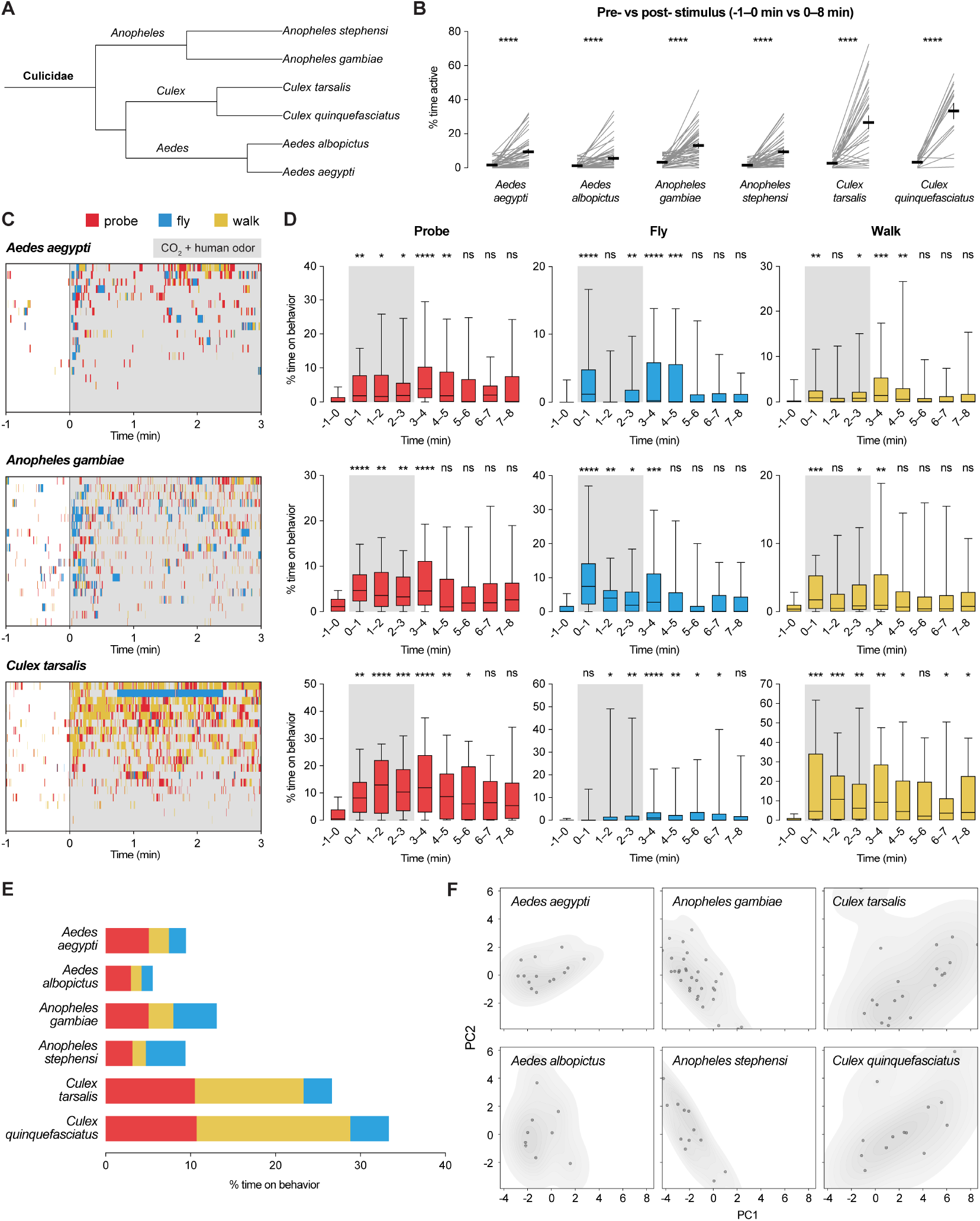
Host-seeking behavioral repertoire varies by mosquito species. (A) Mosquito species tested in this study. Phylogenetic tree generated with Interactive Tree of Life (iTOL)^47^. (B) Percentage of time active pre- vs. post-stimulus (N = 15 – 55 females). Mean and SEM. Individual lines represent single females. Paired t-test. (C) Individual ethograms of 20 randomly-selected individuals coded by behavioral classification (D) and percent of time spent on each behavior (N = 30 – 47 females). The gray shaded bar in group activity plots represents the period of host cue stimulation. Box: median and IQR, whiskers: 5–95 percentile. Kruskal-Wallis test with Dunn’s multiple comparisons test using −1 – 0 min as control. (E) Overall percent of activity in each category during and after stimulus delivery (0 – 8 min) (remaining time spent at rest). (F) PCA plot of host-seeking behavioral repertoire of females that responded during stimulus delivery (0 – 3 min) (N = 10 – 33 females). *P<0.05; **P<0.01; ***P < 0.001; ****P < 0.0001; P > 0.05: not significant (ns). Exact P values are provided in Table S1.

**Figure S2.**
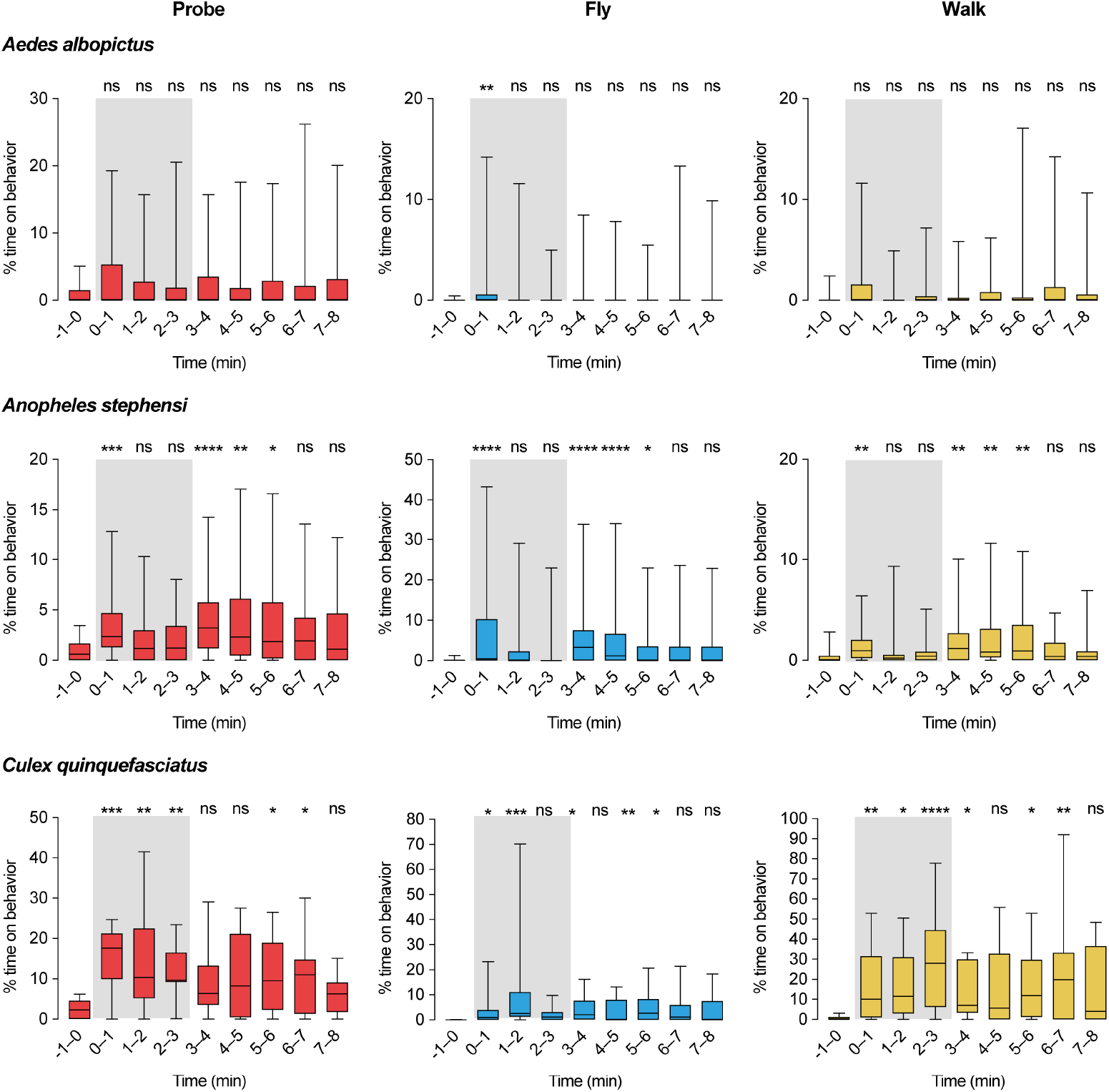
Analysis of host-seeking behavior in *Aedes albopictus, Anopheles stephensi*, and *Culex quinquefasciatus*. Percent of time spent on each behavior (N = 15 – 55 females). The gray shaded bar in group activity plots represents the period of host cue stimulation. Box: median and IQR, whiskers: 5–95 percentile. Kruskal-Wallis test with Dunn’s multiple comparisons test using −1 – 0 min as control. *P<0.05; **P<0.01; ***P < 0.001; ****P < 0.0001; P > 0.05: not significant (ns). Exact P values are provided in Table S1.

### Heterogeneity in host-seeking suppression across mosquito species

We next allowed females to blood feed to repletion and then assessed their behavioral responses to host cues 24, 48, and 72 hours after a blood meal. The timeline of host-seeking suppression of is most extensively characterized in *Aedes aegypti* and our automated behavioral analysis pipeline successfully recapitulated a state of host-seeking suppression, indicated by a loss of host cue-induced locomotion (Figure S3A) and a marked reduction in probing behavior (Figure S3B). This approach provides a high-resolution, temporally dynamic readout of host-seeking behavior that does not rely on prolonged search behavior, required in traditional olfactometer assays. We then applied this analysis to evaluate postprandial host-seeking suppression in all six species. In each case, females consumed meals that approximately doubled their body weights (Figure S4). Although all species exhibited suppressed activity after feeding, the timing and duration of this suppression varied substantially between species.

**Figure S3:**
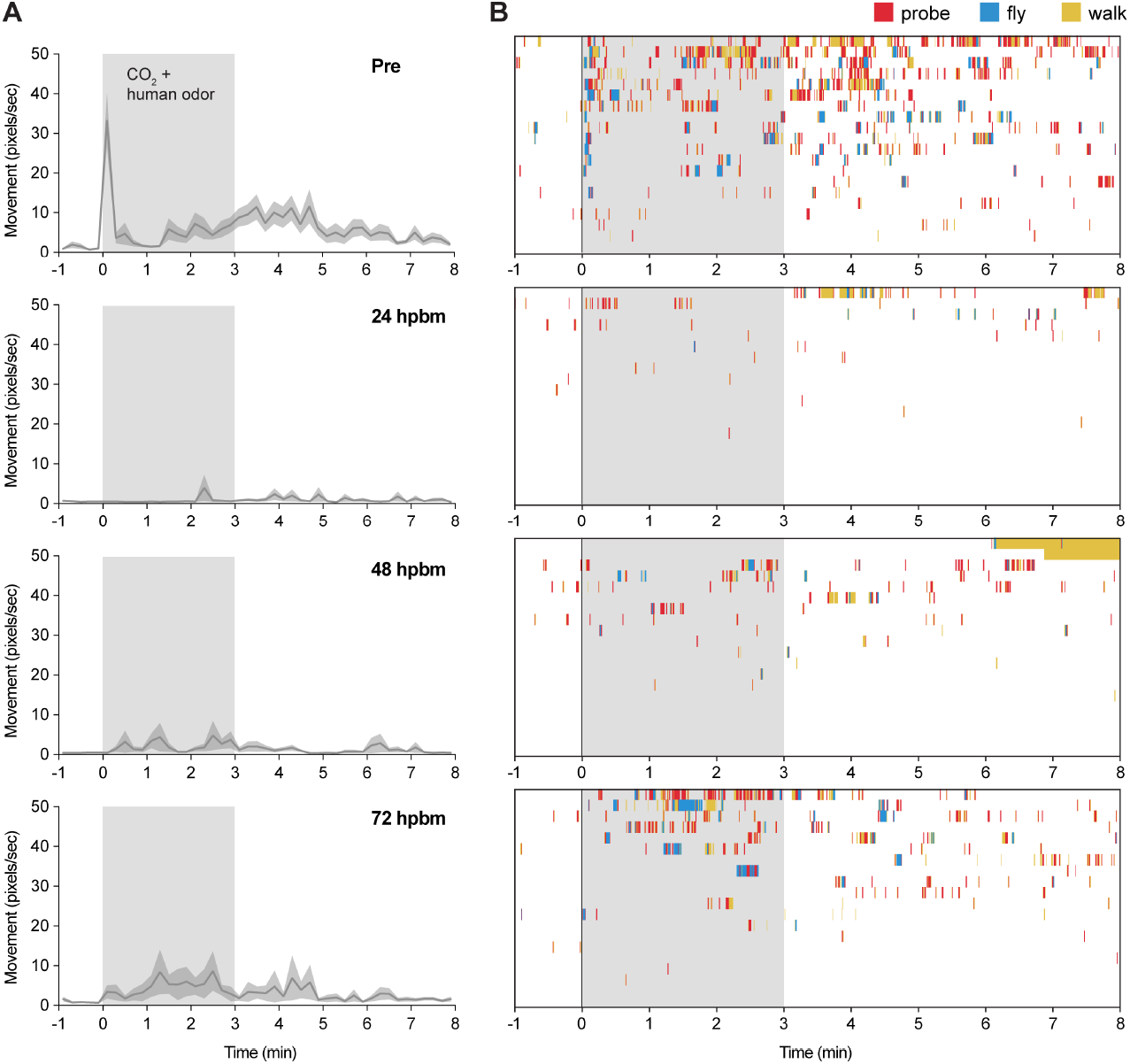
Detailed activity analysis of host-seeking suppression in *Aedes aegypti*. (A) Average movement of females in response to host cues. (N = 21 – 43 females). Mean and SEM. (B) Activity traces of individual females after blood feeding. (20 representative actograms were randomly selected for visual clarity).

**Figure S4:**
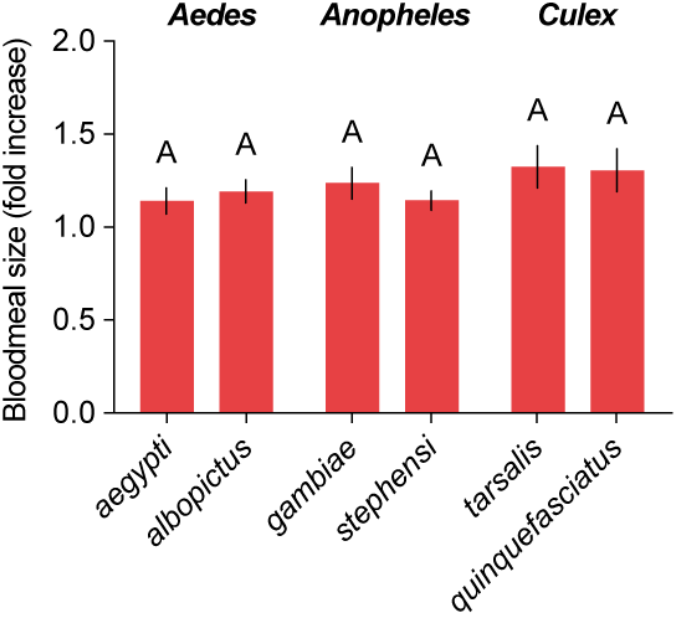
Size of blood-meals ingested by female mosquitoes using an artificial blood-feeder. (N = 20 – 37 females). Data is shown as fold increase in body weight. Mean and SEM. Ordinary one-way ANOVA with Tukey’s multiple comparisons test. Columns are not significantly different from each other P > 0.05. Exact P values are provided in Table S1.

To differentiate general activity from host-seeking behavior, we evaluated both overall movement and probing, which is an indicator of active host-seeking. *Aedes aegypti* females showed an overall suppression of movement and probing with minimal levels at 24 hours that gradually recover to intermediate levels at 72 hours (Figure 3A). In contrast, *Aedes albopictus* females displayed a recovery pattern with levels of movement and probing reaching levels comparable to pre-feeding baseline by 72 hours (Figure 3B). *Anopheles gambiae* females showed prolonged suppression with low levels of movement and probing observed across all timepoints up to 72 hours post-feeding (Figure 3C). In contrast, *Anopheles stephensi* females followed a recovery of probing and movement at 72 hours comparable to the non-fed state (Figure 3D).^48^ *Culex tarsalis* females exhibited a delayed suppression profile, with moderate movement and probing at 24 hours post-feeding followed by a decline at 48 and 72 hours (Figure 3E). This suggests that the abdominal distension from blood meal ingestion might not be sufficient to suppress host-seeking in the short term. In contrast, *Culex quinquefasciatus* females showed significant reduction in movement and probing 24 hours after feeding which was followed by full recovery of movement by 72 hours, though probing remained at intermediate levels (Figure 3F). Overall, at 24 hours post-feeding, all species except for *Culex tarsalis* displayed maximal suppression of both movement and probing (Figure 3G). By 72 hours, several species showed partial or full recovery (Figure 3H).

**Figure 3:**
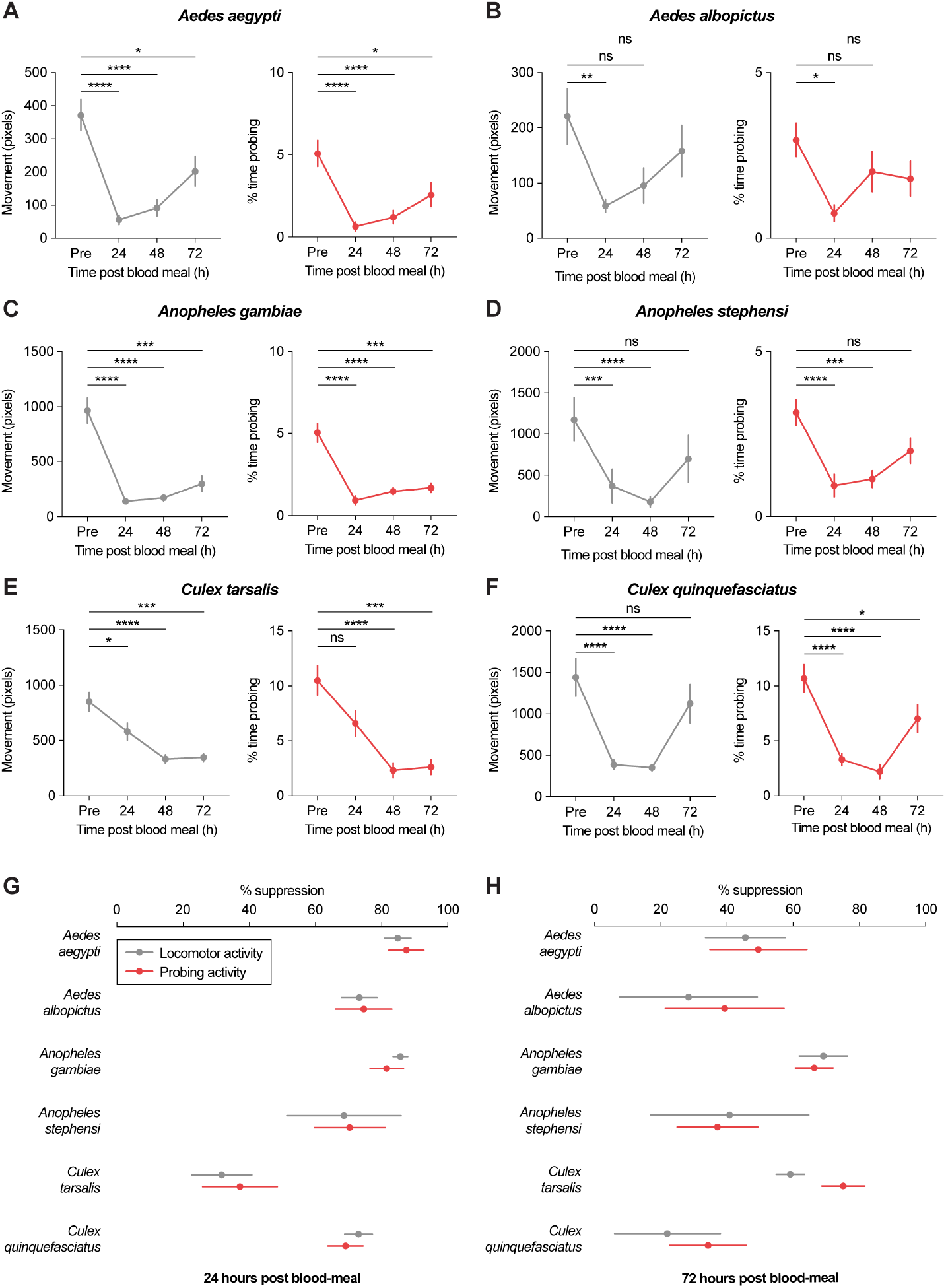
Host-seeking suppression onset and duration vary between species. (A–F) Total movement and % of time spent probing during and post stimulus (0–8 min) in *Aedes aegypti* (A), *Aedes albopictus* (B), *Anopheles gambiae* (C), *Anopheles stephensi* (D), *Culex tarsalis* (E), and *Culex quinquefasciatus* (F) (n = 13–55 females across all species and timepoints). Mean and SEM. Mann-Whitney test. *P<0.05; **P<0.01; ***P < 0.001; ****P < 0.0001; P > 0.05: not significant (ns). Exact P values are provided in Table S1. (G and H) Percent suppression at 24 hours (G) (n = 19–27 females) and 72 hours (H) (n = 13–23 females) post blood meal. Mean and SEM.

## Discussion

With over 120 hours of behavioral tracking and classification data from over 800 individuals across six species, our study established a pipeline for high-temporal-resolution quantification of host-seeking behavior, revealing significant variation in the repertoire of host responsiveness as well as the timeline and extent of host-seeking suppression. The individual arenas were designed to optimize high-quality tracking and allow multiple parallel experiments, ensuring well-matched controls. However, this setup was not optimized for long-range flight, limiting our ability to assess directional navigation. Additionally, we used human odor and CO_2_ as stimuli, which are optimal for species such as *Aedes aegypti* and *Anopheles gambiae*^3,49–51^ but may be suboptimal for *Culex* species, which are more opportunistic and often seek avian hosts.^52^ However, we note that our experimental setup can be easily adapted to test other host odors, temperature cues, or alternative attractants.

Our analysis revealed heterogeneity in the movement repertoire of host-seeking females and of host-seeking regulation after a blood meal. While all species examined exhibited host-seeking suppression after blood feeding, the onset, duration, and intensity of suppression varied. This suppression is believed to have evolved as an adaptive strategy to optimize reproductive success by reallocating energy from host-seeking toward egg development.^37,39^ By temporarily ceasing host-seeking behavior, female mosquitoes can also avoid potentially dangerous host defensive behaviors during this critical period. Interestingly, we observed differences in host-seeking suppression even among closely-related species: *Anopheles gambiae* exhibited prolonged suppression, while *Anopheles stephensi* showed a significant recovery of movement and probing by 72 hours post-feeding. This contrasts with previous studies suggesting that *Anopheles gambiae* females rapidly resume host-seeking^13^, and emphasizes that host-seeking suppression patterns cannot be generalized across an entire genus.

The mechanisms underlying host-seeking suppression may vary between species due to physiological and ecological differences. Larval nutrition can influence host-seeking and refeeding, *Anopheles* mosquitoes are thought to return to host-seeking more quickly because their larval nutritional reserves are lower compared to *Aedes* and *Culex* species.^53^ Abdominal distension plays a key role in suppression, but the threshold for triggering this response may vary between species and be influenced by body size. The delayed suppression observed in *Culex tarsalis* suggests that early cues, such as abdominal expansion, may be processed differently and not directly lead to immediate host-seeking suppression in this species. Neuropeptide signaling also contributes to species-specific differences in host-seeking suppression. Previous studies have identified key neuropeptide signaling pathways involved in *Aedes aegypti* and *Anopheles stephensi*, but suggest that these signals are expressed in different tissues and may function differently.^27–30^ Additionally, variations in the gonotrophic cycle, the time from blood-feeding to oviposition, may influence the duration of suppression. In *Anopheles*, oviposition generally occurs 48-60 hours after blood feeding^12^ whereas oviposition may require more time in *Aedes* and these females can retain eggs and maintain host-seeking suppression if no suitable oviposition sites are available.^54,55^ Notably, *Culex* females may require more than 5 days to oviposit after blood feeding^56^ and these females exhibited high movement relative to probing at the 72-hour timepoint (Figure 3H) suggesting that behavioral activation may represent a non-host-seeking state, potentially for oviposition. Although the timing of egg laying depends heavily on the availability of oviposition sites and female nutrition, in the laboratory most of these species require at least 3 days for egg development and these recovery patterns are consistent with the possibility of multiple blood meals within a single reproductive cycle. Across the six species tested, *Culex tarsalis* females showed the least suppression 24 hours after feeding, *Aedes albopictus* showed the least suppression 48 hours after feeding, and 72 hours after feeding both *Anopheles stephensi* and *Aedes albopictus* showed evidence of a return to probing and host-seeking.

These findings uncover distinct postural and locomotor responses to host cues across six mosquito species from *Aedes, Anopheles*, and *Culex* genera, along with substantial variation in the timing and magnitude of postprandial host-seeking suppression, even between closely related species. This work lays the groundwork for future studies investigating the molecular and neural mechanisms behind behavioral suppression and how they vary between species. The approach can also be expanded to study responses to different hosts or environmental cues. These species-specific behavioral differences have important implications for vectorial capacity, as host-seeking responses impact successful biting and multiple feedings increase the chances of human contact and pathogen transmission.

## Materials and Methods

### Mosquito strains

The following species and strains were used in this study: *Aedes aegypti* Liverpool, *Aedes albopictus* Foshan, *Anopheles gambiae, Anopheles stephensi, Culex tarsalis* (NR-43026), *Culex quinquefasciatus* (NR-43025) (BEI Resources). *Aedes aegypti* Orlando was used for training data.

### Mosquito rearing

All mosquitoes were maintained and reared at 28°C, 70–80% relative humidity with a 12 h light: 12 h dark schedule. *Aedes* and *Culex* eggs were hatched using 1 L of TetraMin fish food suspension in deionized deoxygenated water (one tablet/L). Larvae were fed with TetraMin fish food tablets one daily as needed. *Anopheles* eggs were hatched in deionized deoxygenated water. Larvae were fed with Koi fish food tablets once daily as needed. Male and female mosquitoes were co-housed and adult female mosquitoes used for experiments were between 7 and 14 days old, with *ad libutum* access to 10% sucrose solution. Females were blood-fed on defibrinated sheep blood (Quad5) with 1 mM ATP at 37°C using Hemotek artificial membrane feeders.

### Behavioral tracking and video acquisition

For the behavioral arena, six independent enclosures were created by attaching 3D-printed lids to 6-well tissue culture plates (Celltreat Scientific Products). A piece of mesh was attached to the lid and two holes were also drilled as far apart as possible in each well of the plate to allow for ventilation and to allow airflow of odor stimuli. The boxes for providing odor stimuli were made by cutting acrylic with a laser cutter. These 3d model files and vector files are available at: (https://github.com/Duvall-Lab/UeharaDongDuvall2025).

A modified version of our previously published MozzieDrome assay^57^ was used for recording behavior and supplying odor stimuli. Briefly, the MozzieDrome is a custom-built black acrylic box (60 × 45 × 45 cm) equipped with IR and normal lighting, and an airflow supply system. Airflow was controlled by solenoid valves through 1/4’’ OD tubing and 1/8’’ ID tubing by adjusting the duration of an on/off cycle according to readings from a digital flowmeter. Air containing 10% CO_2_ was supplied to the solenoid valve and passed through a 4.3 x 7.3 x 11.1 cm (inside dimensions) air-tight box (Pelican 1010) containing a human-scented nylon stocking to become odorized. Nylon stockings were worn for 24 hours on the forearm by the same experimenter.

Air and CO_2_ streams were delivered to the behavioral arena via two outlets in a custom-made acrylic odor delivery box (20 × 15 × 3 cm).

A camera mount was installed on a pre-cut camera hole on top of the enclosure and an IR light platform was placed inside of the enclosure to provide illumination for videography. An IR long pass filter was secured inside the enclosure to block visible light and prevent visible light interfering with video acquisition. Back and side IR illumination was provided with a custom-built light box constructed with 1/4’’ clear acrylic sheets covered with two sheets of Kimwipe to provide diffusion.

To minimize the effects of handling, mosquitos were transferred to 6-well plates at least one hour before the experiment and kept undisturbed, avoiding exposure to human odor stimuli. Animals were moved into the MozzieDrome 4 minutes before the start of recording, which included 1 minute pre-exposure, 3 minutes during exposure to 10% CO_2_ and human odor, and 5 minutes post-exposure. To reduce responses to airflow changes, clean air was continuously flowed through the odor delivery box at a rate of 7 SCFH during the entire recording session, including non-stimulus periods. Switching between clean air and odorized air was achieved via computer-controlled solenoid valves without altering total airflow

Behavior was recorded with a machine vision camera (a2A1920-160umBAS, Basler AG, Germany) through the hole at the top of the MozzieDrome and through an IR long pass filter. The frame rate was 60 fps. Behaviors on diurnal (*Aedes aegypti* and *Aedes albopictus*) and nocturnal (*Anopheles gambiae, Anopheles stephensi, Culex tarsalis*, and *Culex quinquefasciatus*) species were recorded in light and dark phase, respectively. We recorded 20 replicates at each time point post blood-feeding (24h, 48h and 72h Post Blood Meal) for each species. Data from non-fed individuals were collected simultaneously with experimental animals housed in alternating wells of the same 6-well plate.

### Key point tracking and behavioral posture classification

The videos were compressed (~30 MB) using ffmpeg (v6.0). Key point tracking analysis was performed using DeepLabCut (DLC, v2.3.5)^45^, and a model for *Aedes* and *Culex* was created using videos of *Aedes aegypti* Liverpool (LVP), *Aedes aegypti* Orlando (ORL) and *Aedes albopictus* Foshan (FOS). The model for *Anopheles* was created from videos of *Anopheles gambiae* and *Anopheles stephesi*. Still images were extracted from videos of a 6-well plate using DLC, and 16 key-points (proboscis, head, thorax, two segments of each leg and abdominal tip) were manually labeled and used as training data. A total of 120 images, for a total of 720 individuals, were used with 150k iterations to generate the model. The video was cropped to each well and analyzed one well at a time to avoid misidentification of an individual. The extracted positional information of all key points was loaded into VAME (v1.0)^46^ and first converted to egocentric coordinates. The velocities of proboscis, thorax, forelegs and hindlegs were added, resulting in 36-dimensional data, which VAME reconstructed into 30 behavioral categories. The classified behavioral poses were visually inspected and manually annotated, then grouped into walk, flight, and probe pose categories.

We first applied a genus-specific noise filter to remove pose fragments with durations in the lowest 10 percentile. Poses from individual frames were then processed to exclude individuals showing > 300 frames of activity (flight, walk, or probe) before stimulus onset (−1 - 0 min). Poses were then transformed into behavioral states by merging gaps < 15 frames if flanked by the same behavioral type. Probe states are further defined to include walking poses that last for less than 120 frames and are directly adjacent to the key probing pose to accurately reflect the probe-walk-probe behavioral sequence of a probing female. For PCA analysis of host responses we excluded nonresponding animals, defined as those that showed no activity for >90% of the 0-3 minutes during stimulus delivery. To analyze travel distance, x-y coordinate information of the thoracic points was extracted and boxcar averaging was used for smoothing to calculate the distance traveled. All code is deposited in GitHub (https://github.com/Duvall-Lab/UeharaDongDuvall2025).

### Quantification of blood-meal size

We provided sheep blood meals (Quad Five) via artificial blood feeders (Hemotek) to examine differences in blood intake among species. On the day before feeding, we removed the sugar solution from the rearing cage and provided a moistened cotton roll. Females were collected and weighed individually before feeding. They were then placed together in a plastic cup or rearing cage and given access to blood for 1 hour and weighed individually again to compare their body weights pre- and post-feeding.

## Resource Availability

Further information and requests for resources should be directed to the lead contact, Laura B. Duvall.

### Materials availability

Requests for reagents generated in this work may be directed to the lead contact, Laura B. Duvall.

### Data and code availability

All code is available on Github (https://github.com/Duvall-Lab/UeharaDongDuvall2025).

All raw data is available in associated data file and videos are available at (https://doi.org/10.5281/zenodo.15478199).

Any additional information required to reanalyze the data reported in this paper is available from the lead contact upon request.

## Acknowledgements

We thank Thomas Gabel for assistance with animal husbandry, Lindy McBride, Conor McMeniman for providing *Anopheles* and *Culex* strains. We thank Richard Hormigo and the Advanced Instrumentation group at the Mortimer B. Zuckerman Mind Brain Behavior Institute for support with assay development. We thank members of the Duvall lab for comments and useful discussions on the manuscript. This work was supported by the following grants: NIGMS (R35 GM137888) (LBD), Beckman Young Investigator Award (LBD), Pew Scholar in Biomedical Sciences Award (LBD), Klingenstein-Simons Fellowship Award in Neuroscience (LBD), JSPS Fostering Joint International Research (#21KK0273) (TU).

## Author contributions

TU and LBD designed experiments. TU performed behavioral experiments and TU and LD performed blood feeding and meal quantification. TU and LD designed behavioral assays, wrote code for the behavioral analyses, and analyzed the data. TU, LD, and LBD prepared the figures and wrote the manuscript with feedback and edits from all authors.

## References

1. The U.S. Department of Health and Human Services and the U.S. Centers for Disease Control And Prevention (2024). The National Public Health Strategy to Prevent and Control Vector-Borne Diseases in People.

2. Bowen, M.F. (1991). The sensory physiology of host-seeking behavior in mosquitoes. Annu Rev Entomol 36, 139–158.

3. Takken, W., and Verhulst, N.O. (2013). Host Preferences of Blood-Feeding Mosquitoes. Annu Rev Entomol 58, 433–453. 10.1146/annurev-ento-120811-153618.

4. Taylor, B., and Jones, M.D. (1969). The circadian rhythm of flight activity in the mosquito Aedes aegypti (L.). The phase-setting effects of light-on and light-off. Journal of Experimental Biology 51, 59–70. 10.1242/jeb.51.1.59.

5. Rund, S.S.C., O’Donnell, A.J., Gentile, J.E., and Reece, S.E. (2016). Daily rhythms in mosquitoes and their consequences for malaria transmission. Insects 7, 1–20. 10.3390/insects7020014.

6. Liu, S., Zhou, J., Kong, L., Cai, Y., Liu, H., Xie, Z., Xiao, X., James, A.A., and Chen, X.G. (2022). Clock genes regulate mating activity rhythms in the vector mosquitoes, Aedes albopictus and Culex quinquefasciatus. PLoS Negl Trop Dis 16, 1–21. 10.1371/journal.pntd.0010965.

7. Godsey, M.S., Burkhalter, K., Delorey, M., and Savage, H.M. (2010). Seasonality and time of host-seeking activity of Culex tarsalis and floodwater Aedes in northern Colorado, 2006-2007. J Am Mosq Control Assoc 26, 148–159. 10.2987/09-5966.1.

8. Paramasivan R, Philip Samuel P, and Selvaraj Pandian R (2015). Biting rhythm of vector mosquitoes in a rural ecosystem of south India. Int J Mosq Res 2, 106–113.

9. Davis, E.E. (1984). Development of lactic acid-receptor sensitivity and host-seeking behaviour in newly emerged female Aedes aegypti mosquitoes. J Insect Physiol 30, 211–215. 10.1016/0022-1910(84)90005-2.

10. Judson CL (1967). Feeding and oviposition behavior in the mosquito Aedes aegypti (L.).I. Preliminary studies of physiological control mechanisms. Biol Bull 133, 369–377. 10.2307/1539832.

11. Davis, E.E. (1984). Regulation of sensitivity in the peripheral chemoreceptor systems for host-seeking behavior by a haemolymph-borne factor in Aedes aegypti. J Insect Physiol 30, 179–183. 10.1016/0022-1910(84)90124-0.

12. Briegel, H., and Hörler, E. (1993). Multiple Blood Meals as a Reproductive Strategy in Anopheles (Diptera: Culicidae). J Med Entomol 30, 975–985.

13. Klowden MJ B.H., Klowden, M.J., and Briegel, H. (1994). Mosquito gonotrophic cycle and multiple feeding potential: Contrasts between Anopheles and Aedes (Diptera: Culicidae). J Med Entomol 31, 618–622. 10.1093/jmedent/31.4.618.

14. Farjana, T., and Tuno, N. (2013). Multiple blood feeding and host-seeking behavior in aedes aegypti and aedes albopictus (diptera: Culicidae). J Med Entomol 50, 838–846. 10.1603/ME12146.

15. Mitchell, C.J., and Millian, K.Y. (1981). Continued Host Seeking by Partially Engorged Culex Tarsalis (Diptera: Culicidae) Collected in Nature. J Med Entomol 18, 249–250.

16. Gillies, M.T. (1954). The recognition of age-groups within populations of Anopheles gambiae by the pre-gravid rate and the sporozoite rate. Ann Trop Med Parasitol 48, 58–74.

17. Klowden, M.J., Blackmer, J.L., and Chambers, G.M. (1988). Effects of Larval Nutrition on the Host-Seeking Behavior of Adult Aedes Aegypti Mosquitoes. Jounal of the American Mosquito Control Association 4, 73–75.

18. Takken, W., Van Loon, J.J.A., Adam, W., and Takken W, van Loon JJ A.W. (2001). Inhibition of host-seeking response and olfactory responsiveness in Anopheles gambiae following blood feeding. J Insect Physiol 47, 303–310. 10.1016/S0022-1910(00)00107-4.

19. Gangoso, L., Aragonés, D., Martínez-de la Puente, J., Lucientes, J., Delacour-Estrella, S., Estrada Peña, R., Montalvo, T., Bueno-Marí, R., Bravo-Barriga, D., Frontera, E., et al. (2020). Determinants of the current and future distribution of the West Nile virus mosquito vector Culex pipiens in Spain. Environ Res 188, 1–11. 10.1016/j.envres.2020.109837.

20. Roth, D., Henry, B., Mak, S., Fraser, M., Taylor, M., Li, M., Cooper, K., Furnell, A., Wong, Q., and Morshed, M. (2010). West Nile Virus range expansion into British Columbia. Emerg Infect Dis 16, 1251–1258. 10.3201/eid1608.100483.

21. Mitchell, C.J. (1981). Diapause Termination, Gonoactivity, and Differentiation of Host-Seeking Behavior from Blood-Feeding Behavior in Hibernating Culex Tarsalis (Diptera: Culicidae). J Med Entomol 18, 386–394.

22. Galun, R. (1963). Feeding Response in Aedes aegypti: Stimulation by Adenosine Triphosphate. Science (1979) 142, 1674–1675. 10.1126/science.142.3600.1674.

23. Klowden, M.J., and Lea, A.O. (1979). Abdominal distention terminates subsequent host-seeking behaviour of Aedes aegypti following a blood meal. J Insect Physiol 25, 583–585. 10.1016/0022-1910(79)90073-8.

24. Klowden, M.J., and Lea, A.O. (1978). Blood Meal Size as a Factor Affecting Continued Host-Seeking by Aedes Aegypti (L.). Am J Trop Med Hyg 27, 827–831. 10.4269/ajtmh.1978.27.827.

25. Klowden, M.J. (1987). Distension-Mediated Egg Maturation in the Mosquito, Aedes aegypti. J Insect Physiol 33, 83–87.

26. Chambers, G., and Klowden, M. (1996). Distention and Sugar Feeding Induce Autogenous Egg Development by the Asian Tiger Mosquito (Diptera: Culicidae). Entomological Society of America 33, 1–7.

27. Brown, M.R., Klowden, M.J., Crim, J.W., Young, L., Shrouder, L.A., and Lea, A.O. (1994). Endogenous Regulation of Mosquito Host-seeking Behavior by a Neuropeptide. J Insect Physiol 40, 399–406. 10.1016/0022-1910(94)90158-9.

28. Duvall, L.B., Ramos-Espiritu, L., Barsoum, K.E., Glickman, J.F., and Vosshall, L.B. (2019). Small-Molecule Agonists of Ae. aegypti Neuropeptide Y Receptor Block Mosquito Biting. Cell 176, 687-701.e5. 10.1016/j.cell.2018.12.004.

29. Dou, X., Chen, K., Brown, M.R., and Strand, M.R. (2024). Reciprocal interactions between neuropeptide F and RYamide regulate host attraction in the mosquito Aedes aegypti. Proc Natl Acad Sci U S A 121, 1–11. 10.1073/pnas.2408072121.

30. Bansal, P., Pillai, R., Babu, P.D., and Sen, S.Q. (2024). Two neuropeptides that promote blood-feeding in Anopheles stephensi mosquitoes. bioRxiv. 10.1101/2024.05.15.594342.

31. Beach, R. (1979). Mosquitoes: Biting Behavior Inhibited by Ecdysone. Science (1979) 205, 829–831.

32. Dittmer, J., Alafndi, A., and Gabrieli, P. (2019). Fat body–specific vitellogenin expression regulates host-seeking behaviour in the mosquito Aedes albopictus. PLoS Biol 17, 1–26. 10.1371/journal.pbio.3000238.

33. Klowden, M.J. (1990). The Endogenous Regulation of Mosquito Reproductive Behavior. Experientia 46, 660–670. 10.1007/BF01939928.

34. Liesch, J., Bellani, L.L., and Vosshall, L.B. (2013). Functional and Genetic Characterization of Neuropeptide Y-Like Receptors in Aedes aegypti. PLoS Negl Trop Dis 7, e2486. 10.1371/journal.pntd.0002486.

35. Castillo, J.S., Bellantuono, A.J., and DeGennaro, M. (2023). Quantifying Mosquito Attraction Using a Uniport Olfactometer. Cold Spring Harb Protoc 2023, 789–794. 10.1101/pdb.prot108175.

36. Jones, M.D.R. (1981). The programming of circadian flight-activity in relation to mating and the gonotrophic cycle in the mosquito, Aedes aegypti. Physiol Entomol 6, 307–313. 10.1111/j.1365-3032.1981.tb00275.x.

37. Klowden, M.J., and Lea, A.O. (1979). Humoral inhibition of host-seeking in Aedes aegypti during oocyte maturation. J Insect Physiol 25, 231–235. 10.1016/0022-1910(79)90048-9.

38. Luxem, K., Sun, J.J., Bradley, S.P., Krishnan, K., Yttri, E., Zimmermann, J., Pereira, T.D., and Laubach, M. (2023). Open-source tools for behavioral video analysis: Setup, methods, and best practices. Elife 12, 1–20. 10.7554/eLife.

39. Klowden, M.J. (1981). Initiation and Termination of Host-Seeking. 27, 799–803. 10.1016/0022-1910(81)90071-8.

40. Sorrells, T.R., Pandey, A., Rosas-Villegas, A., and Vosshall, L.B. (2022). A persistent behavioral state enables sustained predation of humans by mosquitoes. Elife 11, 1–23. 10.7554/eLife.76663.

41. Hol, F.J., Lambrechts, L., and Prakash, M. (2020). BiteOscope, an open platform to study mosquito biting behavior. Elife 9, 1–24. 10.7554/elife.56829.

42. Main, B.J., Marcantonio, M., Spencer Johnston, J., Rasgon, J.L., Titus Brown, C., and Barker, C.M. (2021). Whole-genome assembly of Culex tarsalis. G3: Genes, Genomes, Genetics 11, 1–5. 10.1093/g3journal/jkaa063.

43. Neafsey, D.E., Waterhouse, R.M., Abai, M.R., Aganezov, S.S., Alekseyev, M.A., Allen, J.E., Amon, J., Arcà, B., Arensburger, P., Artemov, G., et al. (2015). Highly evolvable malaria vectors: The genomes of 16 Anopheles mosquitoes. Science (1979) 347, 1–8. 10.1126/science.1258522.

44. Reidenbach, K.R., Cook, S., Bertone, M.A., Harbach, R.E., Wiegmann, B.M., and Besansky, N.J. (2009). Phylogenetic analysis and temporal diversification of mosquitoes (Diptera: Culicidae) based on nuclear genes and morphology. BMC Evol Biol 9, 1–14. 10.1186/1471-2148-9-298.

45. Mathis, A., Mamidanna, P., Cury, K.M., Abe, T., Murthy, V.N., Mathis, M.W., and Bethge, M. (2018). DeepLabCut: markerless pose estimation of user-defined body parts with deep learning. Nat Neurosci 21, 1281–1289. 10.1038/s41593-018-0209-y.

46. Luxem, K., Mocellin, P., Fuhrmann, F., Kürsch, J., Miller, S.R., Palop, J.J., Remy, S., and Bauer, P. (2022). Identifying behavioral structure from deep variational embeddings of animal motion. Commun Biol 5, 1–15. 10.1038/s42003-022-04080-7.

47. Letunic, I., and Bork, P. (2024). Interactive Tree of Life (iTOL) v6: Recent updates to the phylogenetic tree display and annotation tool. Nucleic Acids Res 52, W78–W82. 10.1093/nar/gkae268.

48. Scott, T.W., and Takken, W. (2012). Feeding strategies of anthropophilic mosquitoes result in increased risk of pathogen transmission. Trends Parasitol 28, 114–121. 10.1016/j.pt.2012.01.001.

49. Tuno, N., Kjaerandsen, J., Badu, K., and Kruppa, T. (2010). Blood-Feeding Behavior of Anopheles gambiae and Anopheles melas in Ghana, Western Africa. J. Med. Entomol 47, 28–31.

50. McBride, C.S., Baier, F., Omondi, A.B., Spitzer, S.A., Lutomiah, J., Sang, R., Ignell, R., and Vosshall, L.B. (2014). Evolution of mosquito preference for humans linked to an odorant receptor. Nature 515, 222–227. 10.1038/nature13964.

51. Degennaro, M., McBride, C.S., Seeholzer, L., Nakagawa, T., Dennis, E.J., Goldman, C., Jasinskiene, N., James, A.A., Vosshall, L.B., Degennaro, M., et al. (2013). Orco mutant mosquitoes lose strong preference for humans and are not repelled by volatile DEET. Nature 498, 487–491. 10.1038/nature12206.

52. Rizzoli, A., Bolzoni, L., Chadwick, E.A., Capelli, G., Montarsi, F., Grisenti, M., De La Puente, J.M., Muñoz, J., Figuerola, J., Soriguer, R., et al. (2015). Understanding West Nile virus ecology in Europe: Culex pipiens host feeding preference in a hotspot of virus emergence. Parasit Vectors 8, 1–13. 10.1186/s13071-015-0831-4.

53. Briegel, H. (1990). Metabolic relationship between female body size, reserves, and fecundity of Aedes aegypti. J Insect Physiol 36, 165–172. 10.1016/0022-1910(90)90118-Y.

54. Venkataraman, K., Shai, N., Lakhiani, P., Zylka, S., Zhao, J., Herre, M., Zeng, J., Neal, L.A., Molina, H., Zhao, L., et al. (2023). Two novel, tightly linked, and rapidly evolving genes underlie Aedes aegypti mosquito reproductive resilience during drought. Elife 12, 1–36. 10.7554/eLife.80489.

55. Judson, C. (1968). Physiology of feeding and oviposition behavior in Aedes aegypti (L.) experimental dissociation of feeding and oogenesis. J Med Entomol 5, 21–23.

56. Eastwood, G., Cunningham, A.A., Kramer, L.D., and Goodman, S.J. (2019). The vector ecology of introduced Culex quinquefasciatus populations, and implications for future risk of West Nile virus emergence in the Galápagos archipelago. Med Vet Entomol 33, 44–55. 10.1111/mve.12329.

57. Dong, L., Hormigo, R., Barnett, J.M., Greppi, C., and Duvall, L.B. (2024). Circadian modulation of mosquito host-seeking persistence by Pigment-Dispersing Factor impacts daily biting patterns. BioRxiv. 10.1101/2024.09.19.613886.

